# PCR-Based Detection and Phylogenetic Analysis of *Candidatus Liberibacter asiaticus* in Citrus Orchards Across Nepal

**DOI:** 10.1101/2025.09.19.677271

**Authors:** Richa Giri, Bal Kumari Oliya, Siddartha Gautam, Krishna Das Manandhar

## Abstract

Citrus greening disease, also known as huanglongbing (HLB), is caused by the gram-negative α-proteobacteria *Candidatus Liberibacter* species. This disease poses a significant threat to citrus production worldwide, including in Nepal. This study aimed to conduct the diagnosis and phylogenetic analysis of the citrus greening pathogen in Nepal using both conventional PCR and computational methods. A total of 1,026 samples were collected from thirteen districts across six provinces in the country. PCR-based diagnosis was performed using the primer set Las606/LSS, which targets the 16S rRNA gene of *Candidatus Liberibacter asiaticus*. Additionally, 16S rRNA gene sequencing was performed using sanger sequencing for five samples collected from different geographical regions. The obtained sequences were deposited in GenBank, and a phylogenetic tree was constructed based on these sequences. Among the 1,026 samples tested, 255 were positive, indicating the widespread distribution of HLB across Nepal. All consensus sequences from Nepal showed strong evolutionary relatedness within the *C. L. asiaticus* cluster, displayed over 99% genetic similarity with reference sequences from various parts of the world. Phylogenetic analysis revealed that the Nepalese sequences were closely related to *C. L. asiaticus* sequences from India, such as OM522080.1 (Punjab) and MH473394.1 (Meerut). Additionally, sequences obtained from different regions of Nepal clustered closely together. The findings of this research provide valuable insights into the genetic diversity and prevalence of citrus greening disease, which will aid the development of better management strategies to combat this devastating disease. Furthermore, the sequence data deposited in GenBank will serve as a key reference for future molecular studies on citrus greening disease.

## Introduction

Citrus is grown in tropical and subtropical climates around the world and both fresh citrus and processed citrus products continue to be in high demand globally. It is predominantly produced in China, Brazil and the United States [1]. Mandarin orange is the most extensively cultivated citrus variety in the hilly regions of Nepal, playing a key role in improving food security, nutrition and employment [2]. Citrus varieties such as *Citrus reticulata, C. unshiu, C. sinensis, C. grandis, C. paradisi, C. limon, C. aurantifolia, Fortunella* and *Poncirus trifoliata* are widely cultivated in major regions of Nepal [3]. Mandarin oranges (*C. reticulata*) account for approximately 65% of Nepal’s citrus production, followed by sweet oranges (*C. sinensis*) at 16% and acid limes (*C. aurantifolia*) at 11%, highlighting their dominance in the citrus fruit industry [4]. Citrus fruits alone account for 22.37% of the total fruit production, with cultivation widespread in approximately 60 districts of the country. Despite their dominance in the fruit industry of the country, domestic production is still insufficient to meet the overall demand [5].

Among the different factors of citrus decline, citrus greening is considered the most severe cause of citrus decline worldwide [6]. Citrus greening disease or huanglongbing disease has posed a major threat to the citrus industry worldwide with Nepal being no exception [7,8]. Citrus greening disease was first reported in 1968 and has been rapidly observed in different parts leading to citrus decline in Nepal [9]. Gram-negative bacteria *Candidatus Liberibacter* spp. is the causative agent of citrus greening disease, which is primarily found within the phloem tissues of the plant [10]. The three primary species of *Candidatus Liberibacter* associated with citrus greening disease are *Candidatus Liberibacter asiaticus, Candidatus Liberibacter africanus* and *Candidatus Liberibacter americanus. Candidatus Liberibacter asiaticus* is commonly found in Asia, *Candidatus Liberibacter africanus* is found in Africa, while *Candidatus Liberibacter americanus* is found in South America [11]. There is a limitation in understanding the biology of this pathogen since there is no successful pure (axenic) culture, which limits the functional genomic analysis leading to challenges in HLB disease management [12]. Despite several attempts being made to culture this pathogen only short-term culture has been reported [13]. HLB is a vector-borne disease transmitted by psyllids, also known as jumping plant lice, which feed on phloem sap [14]. *C. L. asiaticus* is transmitted by citrus psyllid vector *Diaphorina citri* Kuwayama and *Candidatus Liberibacter africanus* is transmitted by vector *Trioza erytreae* [4,10]. The primary hosts of *D. citri* are all members of rutaceae family. Major commercial citrus species identified as key hosts include lemon, sour orange, grapefruit, lime, pomelo mandarin and tangerine. Additionally, *Murraya koenigii* (curry tree) and *M. paniculata* (orange jasmine) are also preferred hosts of this vector [15].

Infected trees exhibit several symptoms which can be observed on the leaves, fruit and canopy of the tree [16]. A higher level of the pathogen is found in the petiole, midrib, peduncle and columella while a lower level of the pathogen could be observed in seeds, buds and bark [17]. The distinct yellow pattern on a leaf (blotchy mottle), yellowing of leaf veins, premature leaf drop, poor root system, retarded growth are the major symptoms associated with citrus greening disease [18, 19, 20]. Fruits from infected trees are often deformed, containing abnormal seeds with increased bitterness and sourness with reduced sweetness in the juice [17,19]. Early HLB symptoms can be observed uniformly across the fibrous roots of infected trees [21]. HLB-infected citrus roots show black to dark brown discoloration due to increased polymerization of lignin and tannins. While this lignification acts as a barrier against pathogen invasion, it also reduces the root’s ability to absorb water and nutrients making it prone to water and temperature stress [22]. Whenever the pathogen infects the phloem tissue, they secrete various effectors and virulence factors and also triggers plant immune response which is responsible for callose deposition, cell death and protein accumulation hindering the conductivity of phloem tissue [23].

HLB infects the majority of citrus species, citrus species like grapefruit, sweet oranges, tangelos and mandarins are highly susceptible while lime, lemon and trifoliate are less susceptible compared to other citrus species [19,24]. There is no confirmed transmission of disease through seed [25]. Trifoliate orange has shown low pathogen titer when grafted to infected rootstock making it the least susceptible citrus species [26]. HLB is a graft-transmissible disease [27], side grafts involving twigs are found to be particularly high in pathogen transmission [28]. The grafting method is used to produce citrus saplings in Nepal. Since HLB is highly prevalent, a healthy mother plant is essential to produce citrus saplings. The PCR method is widely used for the diagnosis of HLB prior to the distribution of plants in Nepal [29].

Appropriate and early diagnosis tools are required to prevent the further spread of disease since bacterial load differs variably depending on several factors [30]. Diagnosis of HLB can be done by several methods like electron microscopy, serology, enzymatic assays, polymerase chain reaction (PCR), quantitative PCR (qPCR) [10]. PCR technique is widely used for diagnosis of HLB since it is more sensitive and convenient. Frequently used primers that target 16S rDNA include Las606/LSSS [31] and primer OI1/OI2c [32], primer A2/J5 target the nusG-rplK region [33]. Despite the availability of various primers, PCR amplification is not observed even in severely infected trees [34]. It might be due to several factors like a low pathogen in template DNA, the presence of PCR inhibitors and nonspecific primers [35]. Highly conserved regions of the 16S rRNA gene are used for the construction of universal primers and highly variable regions of 16S rRNA are used for the identification of individual species [36]. Las specific reverse primer LSS (5’-ACC CAA CAT CTA GGT AAA AAC C-3’) and forward primer Las606 (5’-GGA GAG GTG AGT GGA ATT CCG A-3’) is specific to *C. L. asiaticus*. The primer set Las606/LSS was found to be superior to other commonly used primers like A2/J5, OI1/OI2c and MHO353/MHO354. This primer is highly sensitive for conventional PCR, capable of amplifying low template DNA amplifying effectively despite the presence of various contaminants such as ethanol, citric acid, NaCl and sucrose [31].

Phylogenetic analysis is a widely used tool in bioinformatics due to its high reliability and significance [37]. Within a species, phylogenetic analysis aids in unraveling population dynamics, genetic diversity and the evolutionary processes that shape intraspecies variation [38]. The 16S rRNA gene sequence serves as a benchmark for phylogenetic studies. Although being extensively conserved, the existence of interspersed variable regions within the 16S rRNA gene sequence enables the comparison of closely related species [39]. In phylogenetic analysis, the tree is constructed using optimization principles like Maximum Likelihood (ML), Minimum Evolution (ME) and Neighbor Joining (NJ) Method [40]. The NJ method was employed for molecular identification of citrus greening pathogen for Kinnow mandarin in India [41]. The NJ method is known for its high accuracy and minimal assumptions for tree construction and it offers faster computation speeds compared to other methods [42].

Most previous studies have reported the decline of citrus orchards in Nepal [3, 29, 43, 44]. However, these studies did not comprehensively cover all citrus pocket areas of the country and lacked detailed molecular insights. Therefore, the present research aims to detect *C. L. asiaticus* in suspected citrus trees from various citrus-growing regions of Nepal using conventional PCR, followed by Sanger sequencing, to investigate the origin and phylogenetic relationships of the pathogen both within Nepal and in a global context.

## Materials and Methods

### Sample collection

A total of 1,026 citrus leaf samples were analyzed in this study to assess the prevalence of *C. L*. asiaticus, the causal agent of HLB, in Nepal. The study was conducted at the Laboratory of the Warm Temperate Horticulture Centre (WTHC) Kirtipur, Nepal, over two-year period from 2022 to 2024. Samples were methodically collected from various geographical regions throughout Nepal, encompassing several citrus-producing districts. These included Bhojpur in Province 1, Chitwan, Kathmandu, Ramechhap and Sindhuli in Province 3, Gorkha, Lamjung, Myagdi and Syangja in Province 4, Palpa in Province 5, Dailekh and Salyan in Province 6 and Dadeldhura in Province 7. The sampling covered 6 out of the 7 Provinces in Nepal. This broad sampling strategy was designed to provide a representative overview of the HLB distribution across Nepal. To ensure comprehensive disease surveillance, leaf samples were obtained from both screen houses and open orchards, reflecting different cultivation environments and farming practices. The study included a diverse range of citrus plant materials, incorporating both seedlings and grafted plants. These were collected from various citrus varieties to encompass genetic and phenotypic diversity. Among the different citrus varieties analyzed, mandarin (*C. reticulata*) was the most frequently collected sample followed by sweet orange (*C. sinensis*), owing to its status as the most widely cultivated citrus crop in Nepal and its economic significance in both commercial and subsistence agriculture across the country. The study incorporated other citrus varieties, including acid lime, pomelo, unshu mandarin, navel orange and kumquat. It encompassed local citrus types native to Nepal, such as khoku mandarin and banskharka mandarin. Furthermore, newly introduced varieties in the country such as valencia late orange, avana apireno mandarin, imperial mandarin, daisy tangerine and washington navel orange were also included in the study.

### Sample collection and management

The sampling procedure followed the guidelines outlined in HLB-SOP-1, established by the Ministry of Agriculture and Livestock Development, Government of Nepal [45], ensuring a standardized and systematic approach to sample collection. Citrus trees were visually inspected for characteristic symptoms of HLB, such as blotchy mottle patterns and leaf chlorosis. To account for potential variability in bacterial distribution, leaf samples were collected from multiple directions of each tree, with a preference for symptomatic leaves when available. Following the collection, samples were immediately placed in an icebox to maintain sample integrity during transportation to the laboratory. Upon arrival, leaf samples were thoroughly cleaned to remove contaminants and the mid-vein section was excised and cut into small pieces. These mid-vein sections were then frozen in liquid nitrogen and subsequently stored in a deep freezer at −35°C.

### DNA isolation and visualization

DNA isolation was performed using the CTAB method [46] with slight modification to extract high-quality genomic DNA from citrus leaf samples. Approximately 0.2 g of the mid-vein section from each leaf sample was carefully ground into a fine powder using a mortar and pestle in the presence of liquid nitrogen. This step was essential to facilitate cell disruption and ensure the release of cellular contents. Once the tissue was finely ground, it was mixed with 1 ml of CTAB buffer, which was composed of 2% CTAB, 0.5 M EDTA, 5 M NaCl, 1 M Tris-HCl and 0.2% PVP. The resulting mixture was formed into a fine paste, which was then transferred into a 1.5 ml microcentrifuge tube for further processing. The sample was incubated at 65°C in a water bath for 45 minutes to enable the lysis of cells and the release of DNA. During this incubation period, the sample was gently mixed every 10 minutes to ensure uniform cell disruption and complete DNA release. After the incubation step, the sample was centrifuged at 12,000 rpm for 8 minutes to pellet any cellular debris. The supernatant, containing the DNA, was carefully transferred (approximately 600 µl) into a clean, sterile 2 ml microcentrifuge tube. To remove proteins and other contaminants, an equal volume of Chloroform: Isoamyl alcohol (24:1) was added to the supernatant. The solution was then mixed gently on an orbital shaker for 15 minutes to facilitate phase separation. After mixing, the sample was centrifuged at 13,000 rpm for 5 minutes to separate the phases. The upper aqueous phase, containing the DNA, was carefully transferred (approximately 400 µl) into a sterile 1.5 ml microcentrifuge tube. To precipitate the DNA, 40 µl of 3M sodium acetate was added, followed by 500 µl of ice-cold absolute ethanol. The mixture was then centrifuged at 13,000 rpm for 2 minutes, which led to the formation of a DNA pellet. The supernatant was discarded and the DNA pellet was washed with 500 µl of ice-cold 70% ethanol. The sample was then centrifuged again at 13,000 rpm for 1 minute to remove any residual salts. This washing step was repeated twice to ensure the thorough removal of contaminants, resulting in a high-quality DNA sample. After the final wash, the ethanol was carefully pipetted out and the DNA pellet was air-dried at 37°C for 30 minutes in an incubator. Once dried, the DNA was suspended in 100 µl of TE buffer and stored at 4°C for further use. To confirm the quality of the isolated DNA, it was visualized on a 0.8% agarose gel and the DNA bands were observed using a gel documentation system to verify the integrity and purity of the extracted DNA. This step ensured that the DNA was suitable for downstream applications, including PCR and sequencing.

### HLB diagnosis using conventional Polymerase Chain Reaction (PCR)

The DNA extracted from the mid-vein samples was subsequently utilized in conventional PCR for disease diagnosis. For PCR amplification, the forward primer sequence (Las606) 5′-GGAGAGGTGAGTGGAATTCCGA-3′ and the reverse primer sequence (LSS) 5′-ACCCAACATCTAGGTAAAAACC-3′ were used [31] (Fujikawa and Iwanami 2012). This primer pair is designed to amplify 500 base pair DNA fragments in Huanglongbing positive case. PCR amplification was performed using thermal cycler (TC-312) with a 25-well capacity. Each PCR reaction mixture contained 2 µl of template DNA, 0.3 µl each of forward and reverse primers, 6.5 µl of Master Mix (GoTaq Green Promega Corporation) and 5.9 µl of nuclease-free water, resulting in a total volume of 15 µl. The Master Mix (GoTaq Green) is a premixed solution that includes Taq DNA polymerase, dNTPs, MgCl2 and optimized reaction buffers, making it ideal for efficient DNA amplification. Additionally, the mix contains two dyes (blue and yellow) that allow for real-time visual monitoring of the PCR progress during electrophoresis. After amplification, the PCR products were analyzed by 2% agarose gel electrophoresis to visualize the amplicon. A 100 bp ladder marker (Solis Bio Dyne) was run alongside the samples to determine the size of the PCR products. A sample exhibiting a 500 bp band was considered positive for *Candidatus Liberibacter asiaticus*, confirming the presence of the pathogen in the tested sample.

### Sequencing and phylogenetic analysis

#### Sequencing of positive samples and Initial BLAST Analysis

PCR products from five positive samples, representing different districts of Nepal, were submitted to the Center for Molecular Dynamics Nepal (CMDN) for Sanger sequencing. Both forward and reverse primers were used for sequencing to obtain detailed sequence information for phylogenetic analysis. Ten raw sequences generated from these five samples were submitted to the NCBI database. GenBank accession numbers PP916596–PP916605 were assigned to the raw sequences for further reference and analysis.

#### Consensus sequence generation and phylogenetic analysis

The quality of the sequences was analyzed by examining the chromatogram peaks using SnapGene software. A single consensus sequence was generated by aligning the forward and reverse sequences through AliView software [47]. For this purpose, forward and reverse gene sequences of *C. L. asiaticus* obtained from samples collected in different districts of Nepal Kathmandu, Dadeldhura, Dailekh, Myagdi and Palpa, were merged to form single consensus sequences. Specifically, sequence PP916596 was merged with PP916601, sequence PP916597 with PP916602, sequence PP916598 with PP916603, sequence PP916599 with PP916604 and sequence PP916600 with PP916605 to generate single consensus sequences for each sample, which were then used for subsequent phylogenetic analysis. A phylogenetic tree was constructed using the Neighbor-Joining (NJ) method with 10,000 bootstrap replications and this analysis was performed using MEGA 11 software [48].

## Results

### Sample information

Leaf samples were collected from citrus orchards located in various geographical regions of Nepal (Fig 1). During the field collection, several trees displayed characteristic visual symptoms of citrus greening like yellowing and blotchy mottle pattern on the leaves (Fig 2). In addition, infected trees showed poor growth with small fruit size. Few asymptomatic trees with green leaves were also tested to be HLB positive.

**Fig 1.**
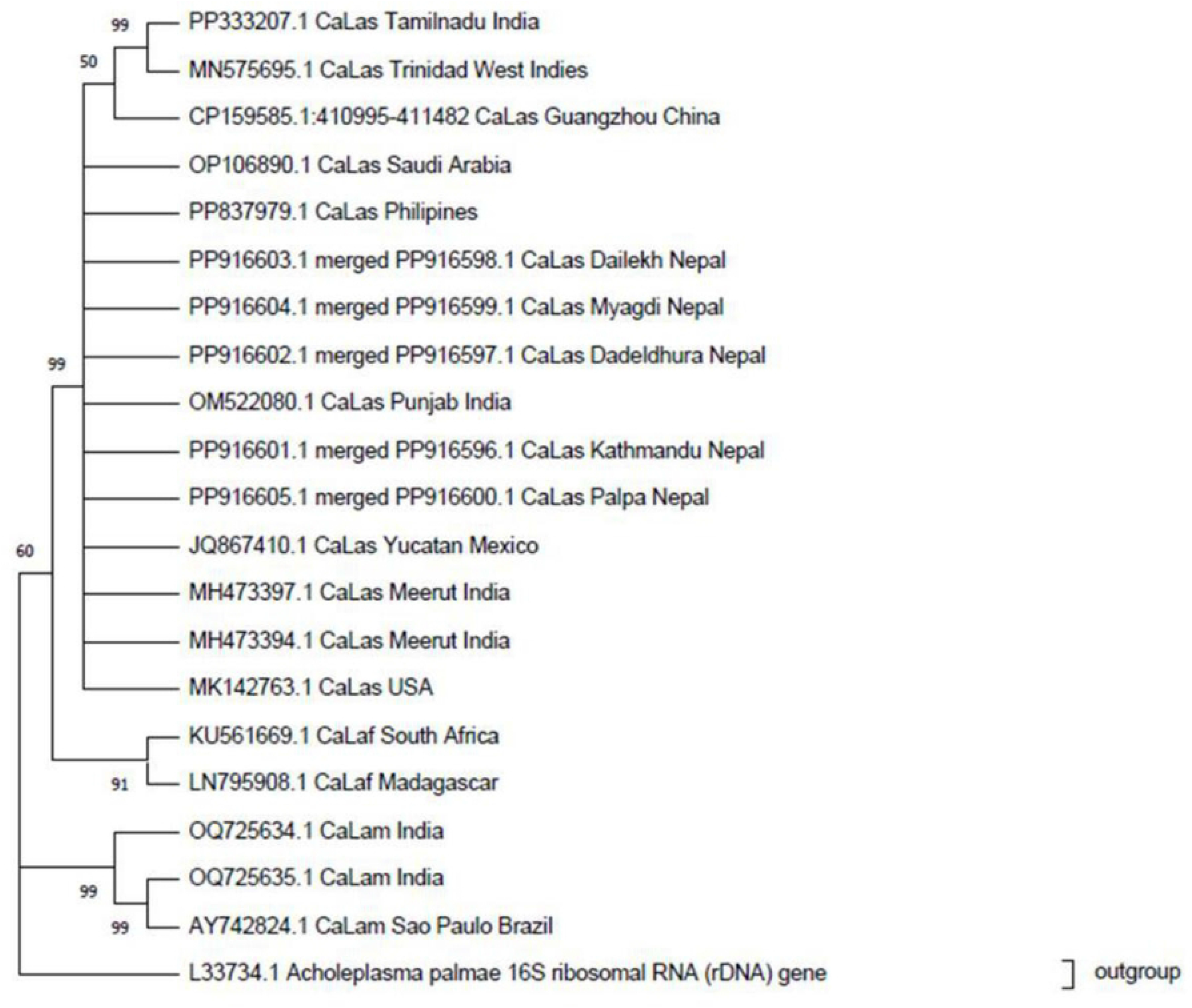
Map of Nepal indicating citrus sample collection sites across different provinces for molecular detection of HLB. Sampling was carried out in selected districts from Provinces 1, 3, 4, 5, 6, and 7.

**Fig 2.**
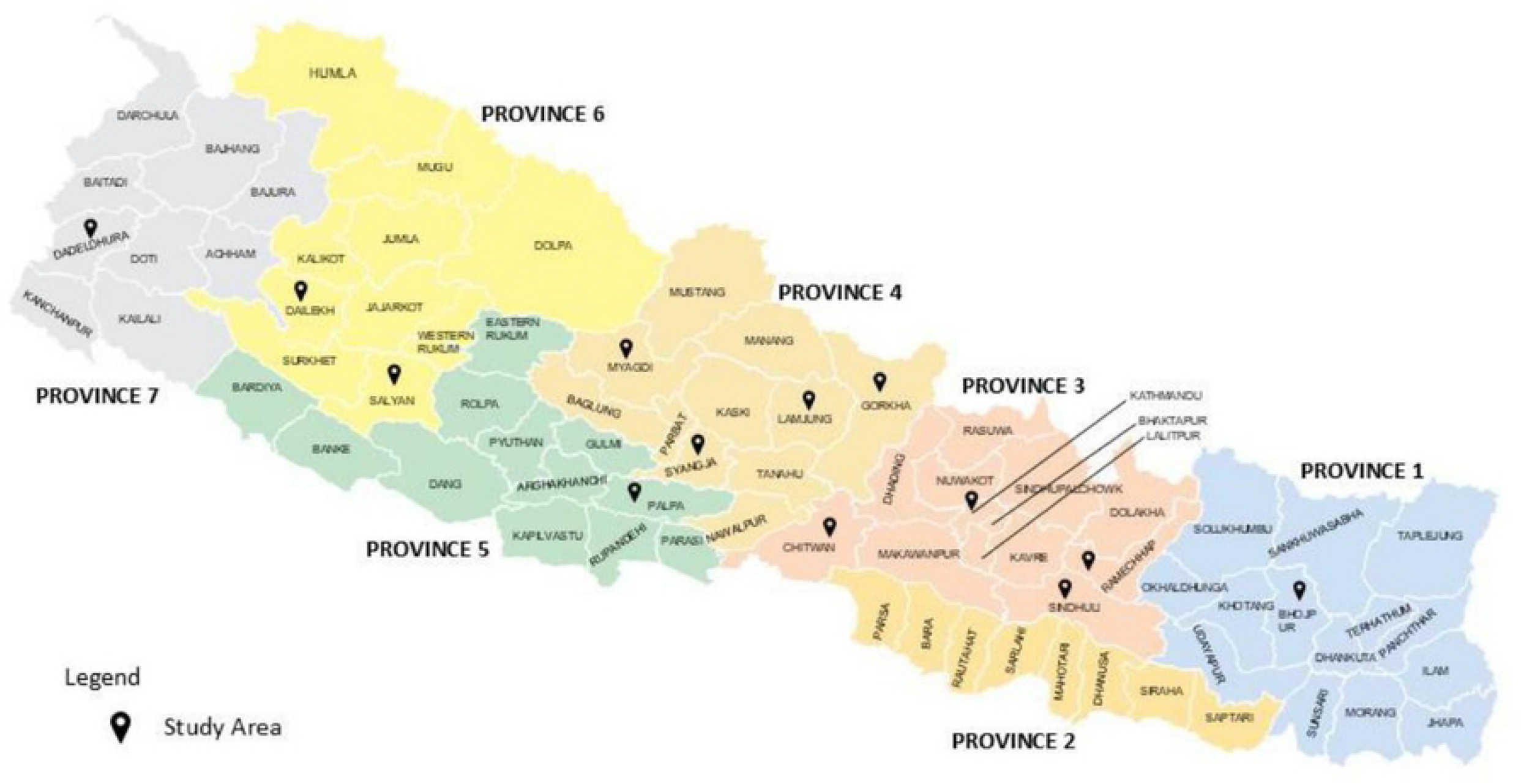
Positive leaf samples exhibiting visual symptoms of citrus greening disease like blotchy mottle and yellowing of leaves.

### DNA extraction and PCR amplification for diagnosis of disease

DNA extraction of all 1026 samples was performed using the CTAB method with slight modifications. After DNA extraction, the quality of the DNA was assessed by visualizing it through gel electrophoresis (Fig 3). Conventional PCR amplification was carried out using the primer set Las606/LSS, targeting the 16S rRNA gene of *Candidatus Liberibacter asiaticus*. Among the 1026 samples collected from various geographical regions of Nepal, 255 samples tested positive for HLB, validated by the presence of a 500 bp amplicon (Fig 4). The result of conventional PCR indicates the prevalence of HLB in multiple areas of Nepal, with significant regional variation. Among the sample tested, the maximum number of positive samples was detected from Ramechhap (74 samples) followed by Gorkha district (43 samples) and Sindhuli (39 samples), sample from Kathmandu and Syangya districts showed low HLB infection (Fig 5). The blue bars represent HLB-positive samples indicating the presence of the citrus greening pathogen, whereas the orange bars represent HLB-negative samples.

**Fig 3.**
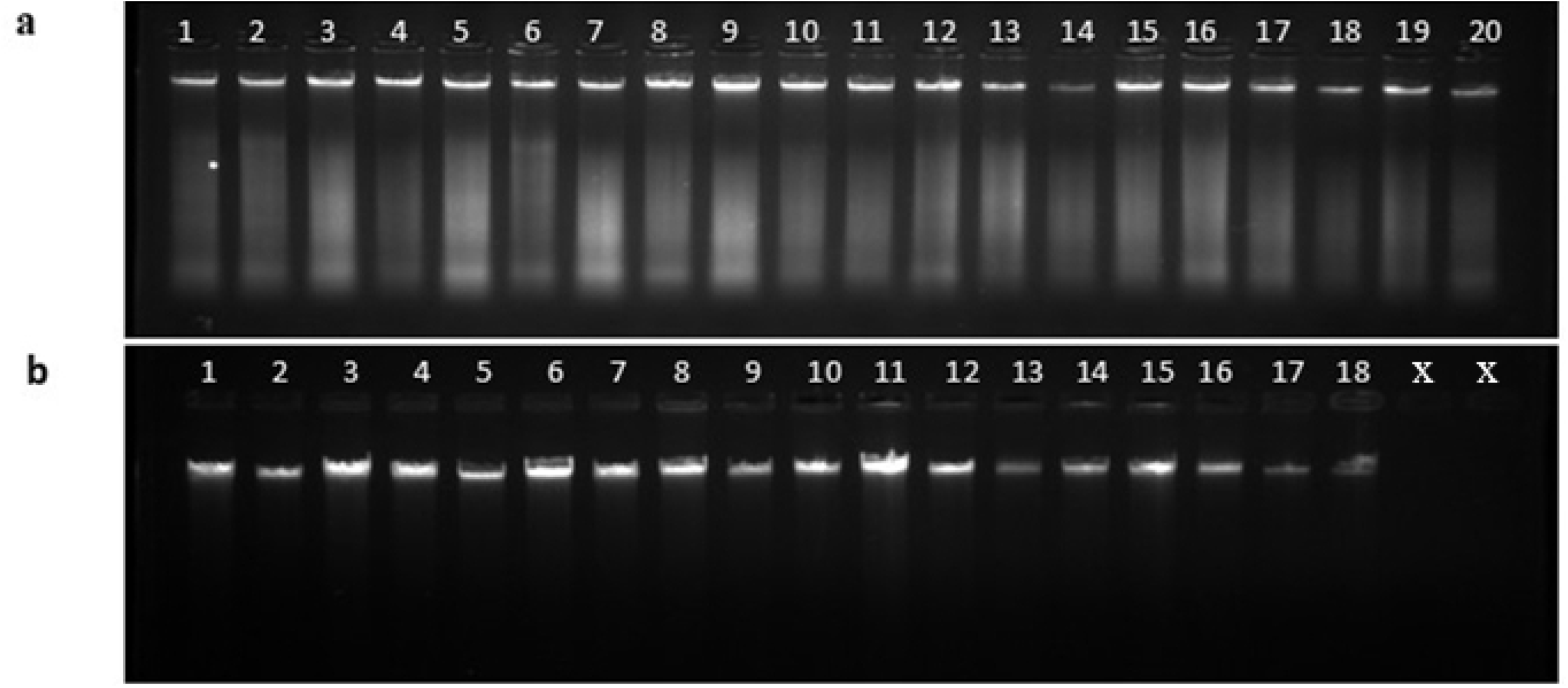
Agarose gel electrophoresis using 0.8% agarose for visualization of DNA extracted from the midvein of citrus leaves collected from various regions of Nepal. (a) DNA extracted from citrus leaf midveins. (b) DNA extracted from citrus leaf midveins from additional samples.

**Fig 4.**
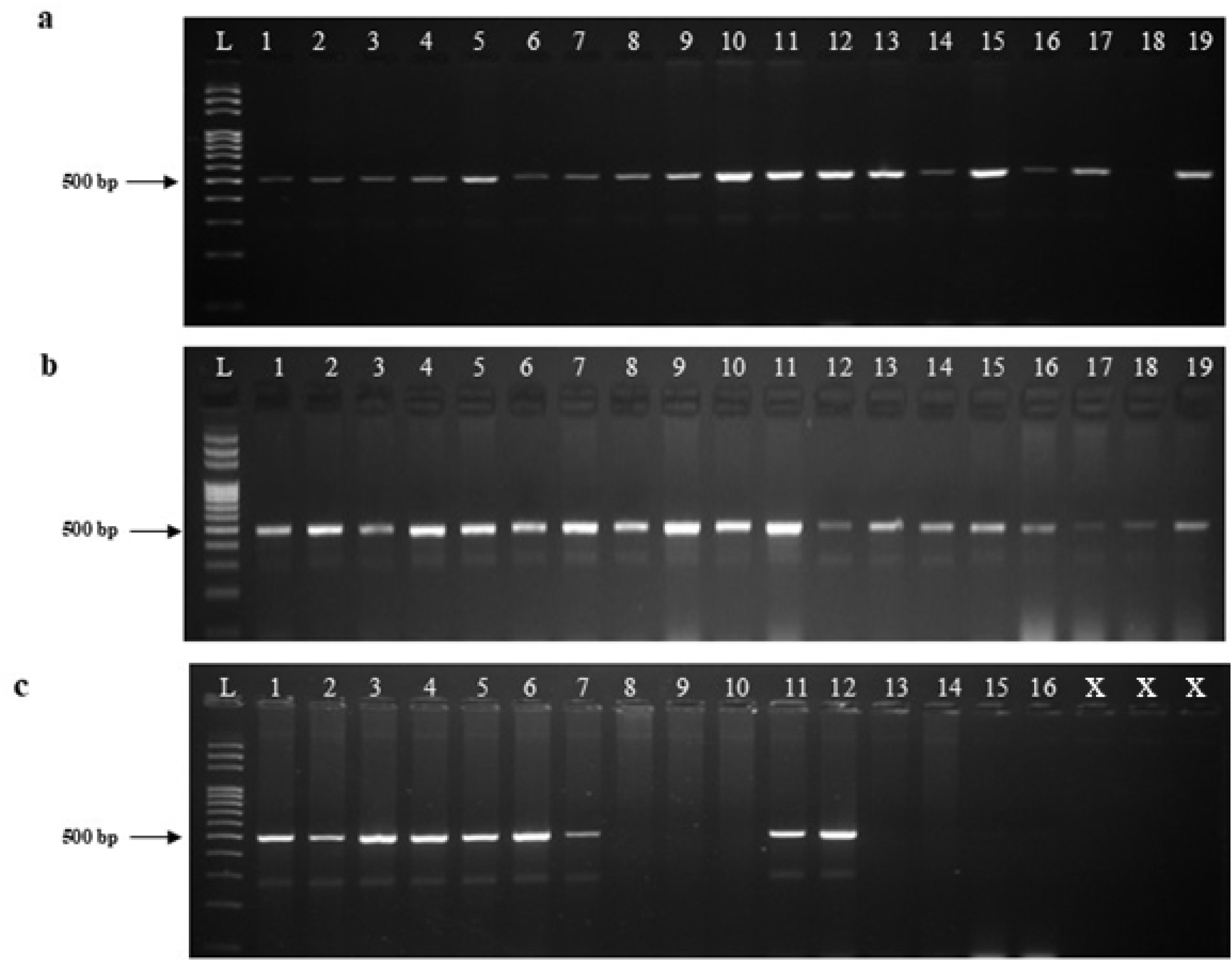
Agarose gel electrophoresis using 2% agarose for visualization of PCR products amplified with the primer set Las606/LSS. Positive samples show a 500 bp amplicon, while negative samples show no amplification. (L) ladder (a) PCR amplification of HLB-positive samples (well 1–17), negative control (well 18), and positive control (well 19). (b) PCR amplification of additional HLB-positive samples (well 1–19). (c) PCR amplification of both HLB-positive and negative samples.

**Fig 5.**
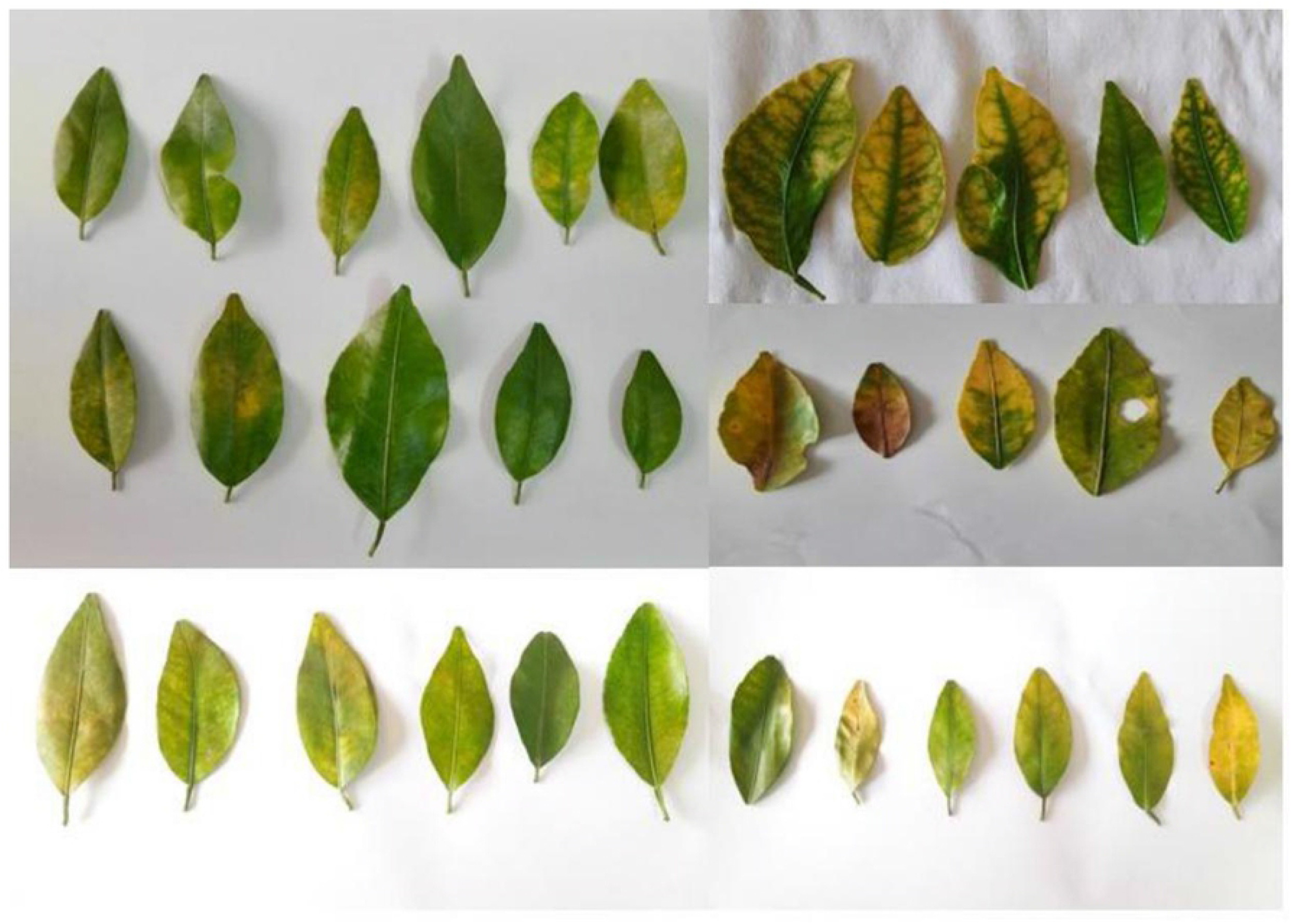
Bar chart showing the number of citrus leaf samples tested for citrus greening disease using PCR across different districts of Nepal. The blue bars represent HLB-positive samples indicating the presence of the citrus greening pathogen, while the orange bars represent HLB-negative samples.

### Sequencing and phylogenetic analysis

The phylogenetic tree was constructed to analyze the genetic relationships of the *C. L. asiaticus* strains identified in the study, based on the sequences obtained from PCR amplification. The Neighbor Joining (NJ) method, a distance-based algorithm, was used to construct the tree. The NJ method calculates the evolutionary distance between different DNA sequences and groups them based on their genetic similarity. All DNA Sequences from Nepal showed strong evolutionary relatedness within the *C. L. asiaticus* cluster, displaying over 99% genetic similarity with the distant sequences from different parts of the world, indicating a possible common origin for these strains. The Basic Local Alignment Search Tool (BLAST) homology search analysis was executed to assess sequence similarity with *Candidatus Liberibacter asiaticus* from various geographical regions. Consensus sequences amplified using Las606/LSS showed greater than 99% homology with *Candidatus Liberibacter asiaticus* 16S rRNA gene sequences: JQ867410.1 (Yucatan, Mexico), OM522080.1 (Punjab, India), OP106890.1 (Saudi Arabia), PP837979.1 (Philippines), CP159585.1 (Guangzhou, China), MH473397.1 (Meerut, India), MH473394.1 (Meerut, India), PP333207.1 (Tamilnadu, India), MK142763.1 (USA), MN575695.1 (Trinidad, West Indies). Additionally, 16S rRNA gene sequences of *Candidatus Liberibacter africanus* (KU561669.1 from South Africa, LN795908.1 from Madagascar) and *Candidatus Liberibacter americanus* (OQ725634.1 and OQ725635.1 from India, AY742824.1 from São Paulo, Brazil) were used to construct a phylogenetic tree. The 16S rRNA gene sequence of *Acholeplasma palmae* (L33734.1) served as an outgroup.

The phylogenetic tree revealed distinct clades of *Candidatus Liberibacter asiaticus* (CaLas), *Candidatus Liberibacter africanus* (CaLaf) and *Candidatus Liberibacter americanus* (CaLam) sequences from various geographic regions and significant branches were supported by high bootstrap values, indicating strong confidence in those groupings. The 16S rRNA gene sequences from different parts of Nepal were clustered in the same clade with *Candidatus Liberibacter asiaticus* sequences obtained from different parts of the world. Within this clade, a subcluster of sequences originating from Tamilnadu, India (PP333207.1), Trinidad, West Indies (MN473394.1) and Guangzhou, China (CP159585.1) were observed with PP333207.1 and MN473394.1 exhibiting a close relationship to one another when compared to CP159585.1. Additionally, 16S rRNA gene sequences of *Candidatus Liberibacter africanus* (KU561669.1 from South Africa and LN795908.1 from Madagascar) and *Candidatus Liberibacter americanus* (OQ725634.1, OQ725635.1 from India and AY742824.1 from Sau Paulo, Brazil) were grouped into separate clades. The CaLaf sequences showed closer evolutionary relatedness to CaLas compared to CaLam, which displayed greater divergence from CaLas (Fig 6)

**Fig 6.**
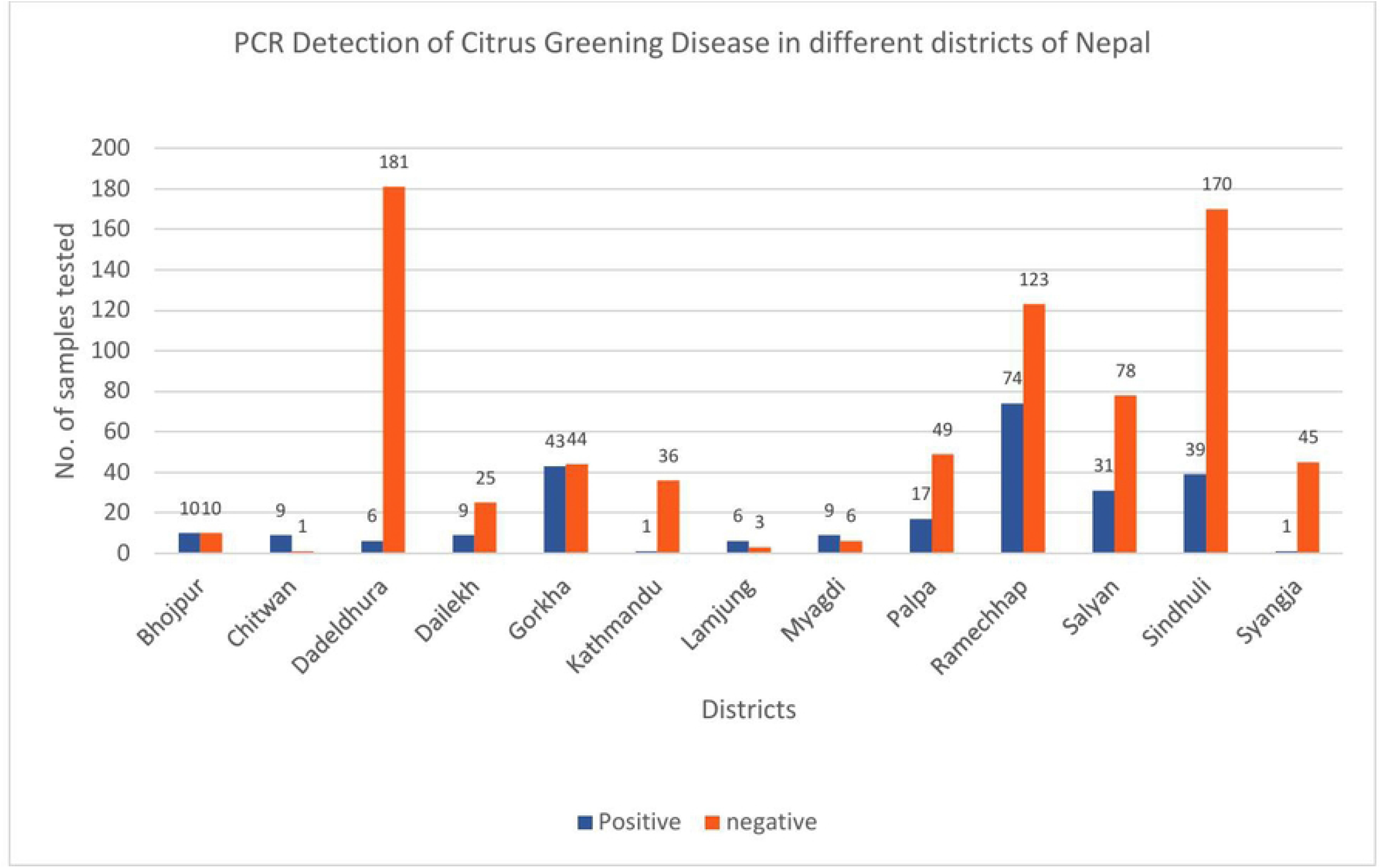
Phylogenetic analysis of 16S rDNA sequences amplified with Las606/LSS primers from five provinces of Nepal and other reported *Candidatus Liberibacter* species using Neighbor-Joining algorithm. The phylogeny was tested using 10000 bootstrap replicates. L33724.1 *Acholeplasma palmale* 16S ribosomal RNA gene was used as an outgroup.

## Discussion

Leaf samples were thoroughly inspected for visual signs and symptoms of HLB before DNA extraction. While some positive samples exhibited the characteristic blotchy mottle pattern and vein yellowing, most leaf samples did not show the classic symptoms of HLB. Instead, these samples displayed symptoms like those caused by nutrient deficiencies. This observation underscores the unreliability of using visual symptoms alone for the diagnosis of HLB, as HLB-positive leaves can exhibit symptoms that overlap with those of other diseases, such as citrus tristeza virus and can also resemble nutrient deficiencies [49]. It is highly unreliable to perform disease diagnosis based only on visual symptoms since the disease might remain asymptomatic due to uneven distribution of bacteria [50, 51]. PCR-based molecular detection offers a more reliable and accurate method for diagnosing citrus greening disease [52, 53].

A total of 1,026 plant samples were analyzed at the Warm Temperate Horticulture Center Laboratory to assess pathogen prevalence across various districts in Nepal. Among these, 255 samples (24.85%) were found to be positive, while 771 samples (75.14%) were found to be negative, indicating a significant variation in disease incidence across different geographical regions. A bar chart was constructed for the number of citrus leaf samples tested for citrus greening disease across different districts of Nepal (Fig 5). The highest positive cases were observed in Ramechhap district (74/197, 37.56%), followed by Gorkha district (43/87, 49.42%) and Sindhuli district (39/209, 18.66%), suggesting these areas as potential hotspots for pathogen transmission. Similarly, Myagdi district (9/15, 60%) and Lamjung district (6/9, 66.67%) also showed high positivity rates; however, the small sample sizes in these districts necessitated further testing to confirm the actual disease prevalence. In contrast, Kathmandu district (1/37, 2.70%), Syangja district (1/46, 2.17%) and Dadeldhura district (6/187, 3.21%) exhibited relatively low positive cases, suggesting either lower pathogen prevalence or successful disease control measures in these areas. Bhojpur district (10/20, 50%) and Chitwan district (9/10, 90%) had high proportions of positive cases, but the small number of samples limited the accuracy of disease prevalence estimation. To gain a more comprehensive understanding of pathogen distribution, it is essential to increase sample size in districts with limited samples, such as Chitwan district, Bhojpur district, Lamjung district and Myagdi district, to ensure statistically significant disease incidence data. The samples for this study were sourced from major citrus-producing regions of Nepal, highlighting a growing concern about the potential decline in citrus orchards due to the spread of HLB. The spread of disease across Nepal may be attributed to grafting with uncertified rootstocks and scions, as well as the improper management of citrus orchards. In an earlier study, Oliya [3] detected *C. L. asiaticus* in various citrus species, except pomelo (*C. grandis*), from Lamjung and Gorkha districts; however, no infection was detected in samples from Dolakha district. In a subsequent study in 2024, Oliya et al. [29] reanalyzed *C. L. asiaticus* in mandarin orange samples from the same orchards in Gorkha, Lamjung, and Dolakha, and found the pathogen present in all three locations. These findings highlight the progressive spread of *Candidatus Liberibacter asiaticus* across different citrus-growing regions of the country. Given that HLB poses a serious threat to citrus production, accurate and reliable detection methods are essential for its effective management and control. In Nepal, conventional PCR remains the primary diagnostic tool for detecting *Candidatus Liberibacter asiaticus*, mainly due to its cost-effectiveness. Although PCR provides high sensitivity and specificity, its effectiveness can be limited by the uneven distribution of the pathogen within infected plants, as the bacteria may not be uniformly present across all tissues [54]. To overcome this limitation, Nepal should consider developing a real-time PCR-based detection system. This advanced technique not only enhances detection sensitivity but also facilitates the identification of the pathogen in both leaf and root tissues. By targeting multiple plant parts, real-time PCR can detect infections even when the pathogen is unevenly distributed, thereby enabling earlier diagnosis and more accurate surveillance of HLB in citrus orchards.

In this study, the diagnosis of all samples was performed using the primer set Las606/LSS, which amplifies a 500 bp segment of the 16S rRNA gene of *C. L. asiaticus*. This primer set is particularly sensitive compared to the previously documented primers [55] (Fujikawa et al. 2013). While other primer sets, such as A2/J5 and OI1/OI2c, are also commonly used for the diagnosis of HLB, the Las606/LSS primer set is preferred for several reasons. One significant advantage of the Las606/LSS primer set is its ability to remain effective even in the presence of contaminants like ethanol and starch, which can inhibit PCR amplification. This makes it a more robust option for field samples that may contain such impurities. Moreover, this primer set is highly sensitive and capable of detecting as low as 1 ng of DNA, ensuring accurate detection even in samples with low concentrations of the pathogen [31]. These characteristics make the Las606/LSS primer set an excellent choice for the molecular detection of *C. L. asiaticus*, offering both reliability and sensitivity in diagnostic applications. HLB-positive samples consistently produced a 500 bp PCR amplicon, which was visualized through agarose gel electrophoresis, confirming the presence of *C. L. asiaticus*. In contrast, no amplification was observed in HLB-negative samples, indicating the absence of the target pathogen. Each PCR assay was carefully designed to include a positive control, which was obtained from a previously confirmed DNA sample, ensuring that the reaction conditions were optimal and the primers were functioning correctly. Additionally, negative control was included in the reaction setup to detect any potential contamination in the PCR master mix or reagents. To enhance the reliability of the results, each PCR reaction was performed in triplicate. Furthermore, in every gel electrophoresis, DNA ladder was used to verify the size of the PCR product.

For the phylogenetic study, a total of five samples representing different geographical regions of Nepal were selected and submitted for Sanger sequencing. Sanger sequencing is widely used in the sequencing field as it offers several prominent advantages: cost-efficiency for sequencing single genes and 99.99% accuracy [56]. DNA sequencing was performed using both forward (Las606) and reverse primer (LSS) sequence. Forward and reverse primers are necessary for Sanger sequencing to ensure accurate and complete sequencing of the target DNA fragment. Moreover, the bidirectional sequencing helps to identify and resolve any potential ambiguities or mutations present in the target sequence. Thus, by comparing the sequences obtained from both direction, high-quality sequencing data can be generated which facilitates accurate interpretation of the results [57].

The evolutionary relatedness of the sequences from different geographical regions can be assessed by analyzing their position within the phylogenetic tree. Sequences obtained from different parts of Nepal were clustered closely suggesting limited genetic diversity within the CaLas populations in Nepal. This might be attributed to the similar environmental conditions from where the samples were taken [58]. CaLas Sequences originating from India, such as OM522080.1 (Punjab) and MH473394.1 (Meerut), grouped relatively close to those from Nepal, indicating a close evolutionary relationship. This suggests that CaLas pathogen may have spread between these regions more recently, possibly through cross-border movement. In contrast, the isolate CP159585.1 from Guangzhou, China, appeared much distant from the sequences from Nepal, highlighting divergence albeit the geographical regions being relatively closer. Similarly, isolates MN575695.1 from Trinidad and MK142763.1 from the USA were considerably distant, reflecting their geographical separation. Additionally, the Philippines isolate PP837979.1 showed moderate relatedness to the sequences from Nepal, implying a shared ancestry within southeast Asia. Although sequences originating from Nepal showed strong evolutionary relatedness to each other, they remained within the broader CaLas cluster while exhibiting strong genetic similarity (greater than 99% sequence similarity) with geographically distant sequences which suggests the existence of a common source or origin for these strains similar to previous study done in India [41]. It is possible that CaLas strains with a common evolutionary lineage were introduced into Nepal through global trade networks or natural dispersal mechanisms. Overall, the close evolutionary relations between CaLas sequences from Nepal and those from around the world highlight the complex dynamics of CaLas transmission and the interconnectedness of citrus-growing regions globally. Understanding these evolutionary relationships is crucial for implementing effective disease management strategies and preventing the further spread of CaLas in Nepal and other citrus-growing areas.

## Conclusions

This study offers critical insights into the detection of Asian citrus greening disease and the genetic relationships among the infected samples. The findings reveal a high prevalence of citrus greening disease across Nepal. Furthermore, phylogenetic analysis based on 16S rDNA sequences demonstrated a close genetic relatedness between *Candidatus Liberibacter asiaticus* strains from Nepal and those from various regions of India, providing valuable information on their evolutionary patterns. These results emphasize the importance of advanced molecular tools for accurate disease diagnosis and genetic characterization of the pathogen. The integration of early diagnostic approaches with phylogenetic studies significantly improves the understanding of the epidemiology of citrus greening in Nepal, ultimately supporting more effective management and control strategies to reduce its impact on citrus production. Additionally, the sequence data deposited in National Center for Biotechnology Information (NCBI) database will serve as a valuable genetic resource for future research on *Candidatus Liberibacter asiaticus*.

## Acknowledgements

We thank the field staff and farmers for their assistance in sample collection and Ms. Sunita Khadka for laboratory support. We acknowledge the Warm Temperate Horticulture Center, National Center of Fruit Development, Ministry of Agriculture and Livestock Development, Government of Nepal, for their support.

## Author’s Contributions

RG: Performed the experiments; Analyzed and interpreted the data, and wrote manuscript; BKO: conceptualized and designed the research, supervised the overall research and critically revised the manuscript, SG: contributed on sequence and phylogenetic analysis, KDM: supervised and critically revised the manuscript.

## Supporting Information Caption

**Supporting Information 1. Sequence data that support the findings of this study have been deposited in the National Center for Biotechnology Information (NCBI) with primary accession code PP916596 to PP916605**

**Supporting Information 2. Nucleotide sequence submitted to the NCBI database: Supporting Information 3. Consensus sequence used in the construction of phylogenetic tree generated using SnapGene and AliView software:**

